# Exploring penetrance of clinically relevant variants in over 800,000 humans from the Genome Aggregation Database

**DOI:** 10.1101/2024.06.12.593113

**Authors:** Sanna Gudmundsson, Moriel Singer-Berk, Sarah L. Stenton, Julia K. Goodrich, Michael W. Wilson, Jonah Einson, Nicholas A Watts, Genome Aggregation Database Consortium, Tuuli Lappalainen, Heidi L. Rehm, Daniel G. MacArthur, Anne O’Donnell-Luria

## Abstract

Incomplete penetrance, or absence of disease phenotype in an individual with a disease-associated variant, is a major challenge in variant interpretation. Studying individuals with apparent incomplete penetrance can shed light on underlying drivers of altered phenotype penetrance. Here, we investigate clinically relevant variants from ClinVar in 807,162 individuals from the Genome Aggregation Database (gnomAD), demonstrating improved representation in gnomAD version 4. We then conduct a comprehensive case-by-case assessment of 734 predicted loss of function variants (pLoF) in 77 genes associated with severe, early-onset, highly penetrant haploinsufficient disease. We identified explanations for the presumed lack of disease manifestation in 701 of the variants (95%). Individuals with unexplained lack of disease manifestation in this set of disorders rarely occur, underscoring the need and power of deep case-by-case assessment presented here to minimize false assignments of disease risk, particularly in unaffected individuals with higher rates of secondary properties that result in rescue.

## INTRODUCTION

Accurately predicting disease risk in asymptomatic individuals, for example in prenatal diagnosis as well as cancer and other later-onset disorders, is critical for realizing precision medicine. Incomplete penetrance, defined as the absence of the disease phenotype in individuals with a disease-associated variant, presents an added challenge in predicting risk and determining variant pathogenicity, especially in yet unaffected individuals. Genome sequencing of apparently healthy individuals frequently identifies presumed disease-causing variants in the absence of reported disease^1–5^. This could suggest that incomplete penetrance is either a surprisingly common feature of human disease or alternatively may be driven by limitations in the accuracy of variant-calling and annotations.

Disease penetrance and its underlying mechanisms are not well understood. Historically, penetrance estimates have relied on symptomatic proband-identified disease cohorts, resulting in overestimating disease risk due to ascertainment bias^6^. Analysis of large-scale population databases and biobanks can provide a less biased avenue to investigate penetrance^7^. Recent penetrance studies utilizing population data have focused on assessing variant penetrance for specific disorders, e.g. prion disease^8^, metabolic conditions^9^, and maturity-onset diabetes of the young (MODY)^10^, and more broadly investigating the association of pathogenic variants to 401 phenotypes in 379,768 individuals from the UK Biobank^5^. Investigations moving beyond descriptive reports and association studies towards focusing on mechanisms underlying incomplete penetrance are few and disease-specific^11–14^, with examples of how expression levels^15^, eQTL^16^, and sQTLs^17^ could modulate penetrance. A statistical approach to investigate eQTL association with incomplete penetrance in neurodevelopmental disease in 1,700 trios from the Deciphering Developmental Disorder cohort did not find altered gene expression due to known eQTLs to be an explanation for incomplete penetrance in the unaffected parents carrying the variant of the affected probands^18^.

The Genome Aggregation Database (gnomAD) is a widely used publicly available collection of population variant data, currently sharing harmonized data from 807,162 individuals, including 76,215 genomes and 730,947 exomes (version 4, released November 2023). The gnomAD dataset has played a key role in supporting the discovery of genes and variants association with genetic diseases, enabling improved variant classification in clinical as well as mechanistic-focused interpretation of variants in disease-discovery research settings^19–21^. Investigation of well-established and predicted disease-associated variants in unaffected individuals in gnomAD presents an unexplored opportunity to improve our variant interpretation abilities, increase understanding of variant effect, and inform on mechanisms affecting the penetrance of disease. The unprecedented size, rigorous quality control pipelines with joint variant-calling over all samples, and diverse ancestry make gnomAD an excellent resource for large-scale analysis of variants reported as pathogenic in clinical databases (e.g., ClinVar). Because phenotype data for individuals in gnomAD is not systematically collected nor able to be shared, we focus on variants associated with severe, dominantly inherited diseases not typically found in individuals recruited for common disease studies or biobanks, from which gnomAD samples originate.

ClinVar is a publicly accessible database that collects submissions mainly from diagnostic laboratories and some research studies, including the clinical significance of variants, and optionally, the associated disease, as well as applied evidence^22^. The open submission model allows the collection of a massive variant classification dataset, which powers large-scale population-based studies; to date, over 3.6 million records have been submitted, and over 2.4 million unique variants have been assigned a clinical significance of pathogenic (P), likely pathogenic (LP), uncertain significance (VUS), likely benign (LB), or benign (B)^22^, with most submitters using recommended standards from the American College of Medical Genetics and Genomics (ACMG) and the Association for Molecular Pathology (AMP)^23^. An inevitable consequence of a crowd-sourced, voluntary, point-in-time submission model is the risk of outdated and/or inaccurate pathogenicity classifications, hence these data must be interpreted cautiously. Having multiple submitters agree on classification and sharing the evidence used towards the classification builds confidence, but 77.3% of variants are currently classified by only one laboratory^24^.

Loss of function (LoF) variants (here including nonsense, essential splice site, and frameshift variants) have important implications in disease biology and are an especially interesting group of variants from an incomplete penetrance perspective as they are considered to have a fairly uniform variant-to-function effect through nonsense-mediated mRNA decay. Thus, any true LoF variant in a haploinsufficient disease-gene is expected to result in disease. In recent work, we provided a protocol for improved evaluation and classification of predicted LoF (pLoF) variants and demonstrated that deeper pLoF assessment beyond standard annotation pipelines is crucial to reduce pLoF false-positive classification rates^25^. The framework presented a set of 32 rules designed based on previously accepted pLoF evasion mechanisms and artifact modes to assess the presence of local modifying variants, the biological relevance of the site, and evidence of a variant being an artifact (a challenge in population datasets where variants have not been analytically validated by an orthogonal method). Each pLoF variant is labeled according to the criteria of these rules, which adds up to a final verdict of LoF, Likely LoF, Uncertain LoF, Likely Not LoF, or Not LoF for each pLoF variant^25^. Further study of pLoF variants associated with disease in presumably unaffected individuals in population databases can enhance our understanding of pLoF functional impact and highlight the possibility of incomplete penetrance in the investigated conditions. Although the vast majority of P/LP variants in any population database are likely to be variants causing disease in an autosomal recessive (AR) manner and any carrier of one allele would simply be a carrier of AR disease, there are also observations of P/LP variants reported to cause disease in an autosomal dominant (AD) manner. Naturally, some of these variants will be expected in common disease studies or biobanks due to being hypomorphic or associated with mild phenotypes, having known incomplete penetrance or variable expressivity, or being late-onset conditions that have not manifested at the time of enrollment, but some are expected to be much less common or even absent in population databases due to the nature of the phenotype (severe, early-onset, highly penetrant).

In this study, we explore disease-associated variants in 807,162 individuals from gnomAD to increase understanding of the underlying reason for tolerance of presumed disease-causing variants and mechanisms of incomplete penetrance. Specifically, we have (1) explored the landscape of ClinVar variants in these individuals, including investigation of how representation varies between gnomAD releases, (2) investigated the prevalence of modifying variants as an explanation for incomplete penetrance in a subset of P/LP variants, (3) deeply investigated all pLoF in 76,215 gnomAD genome variants associated with a set of 77 severe, early-onset, highly penetrant haploinsufficiency disorders for lack of disease manifestation due to limitation in variant annotation, calling somatic variants, detecting artifacts and by mechanisms of incomplete penetrance in truly pathogenic variants. These large-scale analyses of presumed disease-causing variants in the general population provide valuable insight into disease-variant interpretation and the identification of modifying variants that can result in incomplete penetrance.

## RESULTS

### Improved representation of ClinVar variants in gnomAD v4

We investigated the extent to which variants reported in ClinVar are also represented in gnomAD. Of 2,314,231 unique ClinVar variants, filtered to include all single nucleotide variants and indels (<50 base pairs) with assigned clinical significance (P/LP, VUS, B/LB or conflicting classifications), 1,702,421 variants (73.6%) were present in at least one of 807,162 individuals in gnomAD. As expected, we observed a lower representation of P/LP ClinVar variants 66,571/221,975 (30,0%) and a higher representation of B/LB, VUS, and variants with conflicting classifications (73.1%, 83.6%, and 88.8%; Fig. 1a-b). P/LP variants found in gnomAD are of more deleterious variant classes (e.g., pLoF variants: nonsense, frameshift, and essential splice variants), compared to B/LB that are of less deleterious variant classes (e.g., intronic and synonymous variants). VUS mostly consists of missense variants (Fig. 1c). In addition to a lower representation of ClinVar P/LP variants in gnomAD, P/LP variants are rarer compared to other classes. 97.6% of P/LP variants have an allele frequency (AF) of less than 0.01% (AF<0.0001, Fig. 1d, Table S1); 61.1% are observed in five or fewer individuals, and 29.8% in a single individual in gnomAD (Fig. 1e, Table S2). The 66,571 P/LP variants are found on 8,110,001 alleles in 807,162 individuals, resulting in an average of 10.0 pathogenic alleles per individual. Of 66,571 P/LP variants, 63,646 variants could be assigned an inheritance pattern based on the reported inheritance pattern in OMIM. As expected, P/LP variants found in gnomAD are primarily found in genes associated with disorders with an AR inheritance pattern, and there is an underrepresentation of P/LP variants in genes associated with disorders with an AD inheritance pattern compared to variants reported in ClinVar (Fig. 1f).

The 5.7-fold increase from 141,456 to 807,162 individuals from gnomAD v2 to v4 has resulted in improved representation of unique ClinVar variants, from 56.9% in v2 to 73.6% in v4. Representation of P/LP variants has close to doubled and increased from 16.3% to 30.0% of unique P/LP ClinVar variants, and B/LB variants have increased from 70% to 83.6% (Fig. 1a-b, Fig. S1a-b). Reassuringly, distributions of variant types, AF, allele counts (AC), and inheritance are largely consistent between the datasets (Fig. 1c-f, Fig. S1c-f), with a trend towards ClinVar variants having a lower AF as gnomAD population size increases, i.e., an observed lower AF in v4 compared to v2 (Fig. S2). The average number of P/LP variants per individual is similar between versions, 8.8 per individual in gnomAD v2 (1,240,951 alleles in 141,456 individuals) compared to 10.0 per individual in gnomAD v4.

**Fig. 1:**
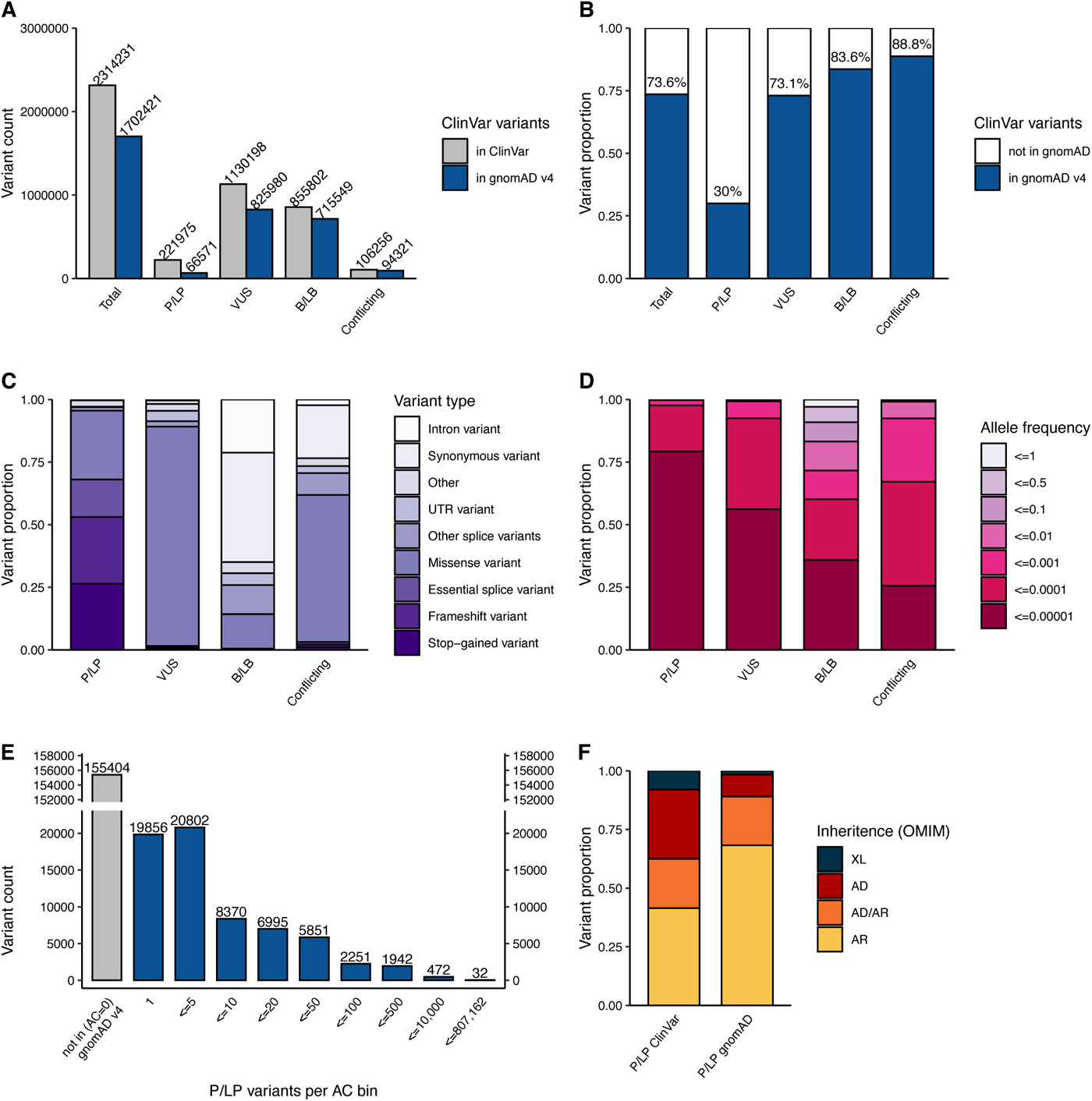
Representation of ClinVar variants in 807,162 individuals in gnomAD v4. (**A**) Variant count of ClinVar variants in ClinVar (grey) vs. gnomAD (blue) in each classification category pathogenic/likely pathogenic (P/LP), variant of uncertain significance (VUS), benign/likely benign (B/LB) or with conflicting classifications. (**B**) Percentage of ClinVar variants reported in gnomAD in at least one individual. (**C**) The proportion of variants by variant type within each clinical significance classification and, (**D**) within each allele frequency bin, both for variants in gnomAD. (E) Total number of P/LP variants within each allele count bin (including variants absent from gnomAD for comparison). (**F**) The inheritance pattern of the gene harboring the P/LP variants in gnomAD versus all variants in ClinVar.

### Example of genetic ancestry group-specific incomplete penetrance

We investigated if a set of 3957 P/LP variants found in 31,014 individuals were tolerated due to modifying pLoF variants in the same gene, potentially reducing the expression of the pathogenic allele. We included all P/LP ClinVar variants in gnomAD with AC ≤50, located in a gene not constrained for LoF (predicted loss-of-function intolerance <0.9, pLI), and with at least one condition of AD inheritance in OMIM, suggesting that pathogenicity is more likely to act through a gain-of-function mechanism.

One example was a pathogenic variant in *GJB2* p.Gly45Glu (13-20189448-C-T) that is reported to cause a severe form of keratitis-ichthyosis-deafness syndrome but found in 35 individuals in gnomAD v4. The condition is lethal due to severe skin lesions, infections, and septicemia^26,27^, through a dominant negative effect resulting in disturbed ion channel transportation. Our analysis confirmed that all 35 individuals also had a downstream nonsense p.Tyr136Ter variant (13-20189174-G-T). Access to paired individual-level sequencing read data from one individual confirmed that the two variants were present *in cis.* This specific p.Tyr136Ter^28^, as well as many other nonsense variants, is associated with AR hearing loss suggesting that the impact of the combined p.Gly45Glu and p.Tyr136Ter allele in these individuals is converted to loss-of-function. The haplotype is likely identical by descent with 34 out of 35 gnomAD individuals belonging to the East Asian genetic ancestry group (Fig. 2). Our analysis suggests that this is a rare example of incomplete penetrance with no evidence for this being a common mechanism in the general population (Supplementary report of results in Note S1, Table S3).

**Fig. 2:**
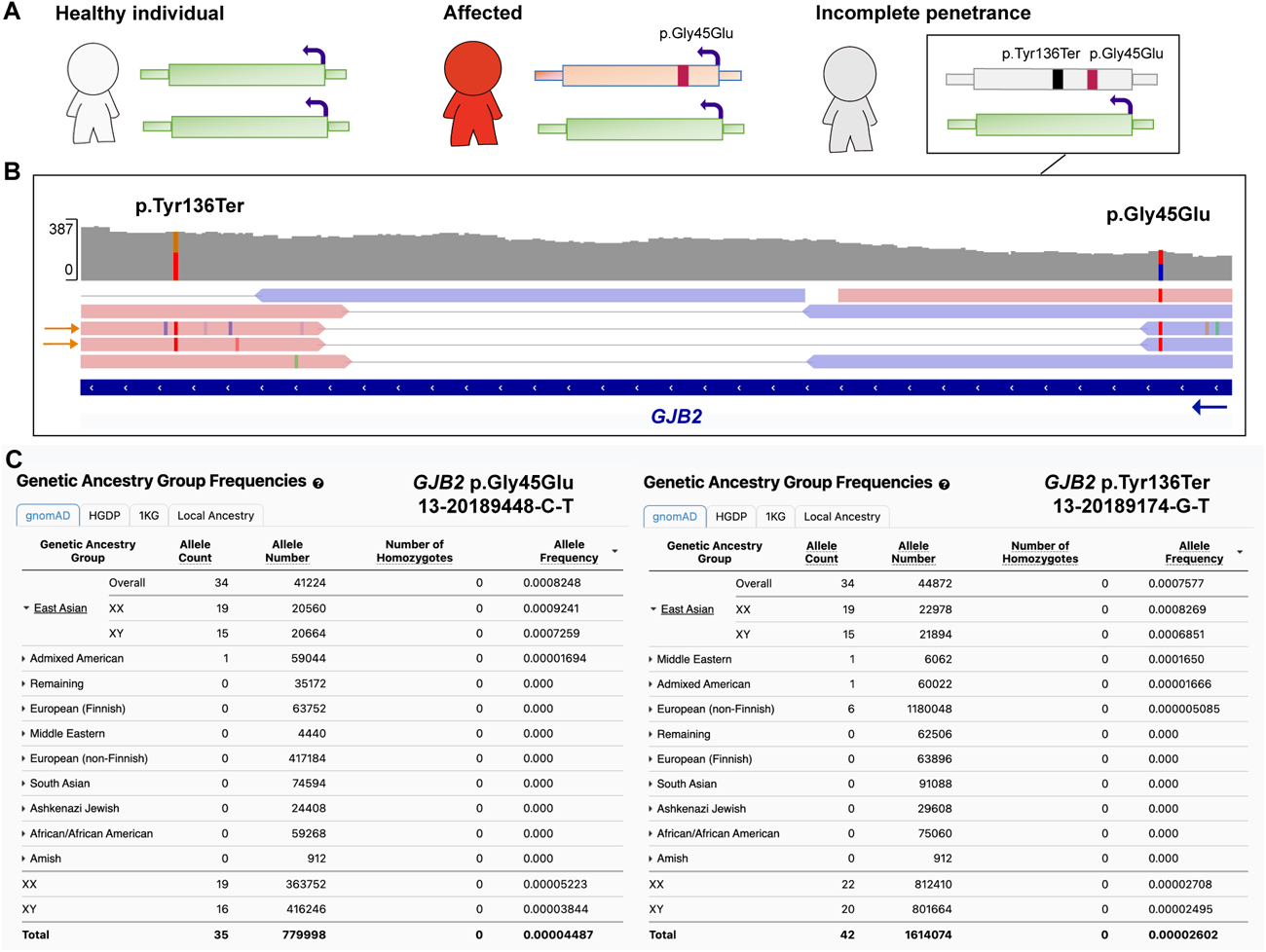
The p.Gly45Glu *GJB2* variant associated with severe pediatric disease is rescued by a modifying variant in the East Asian genetic ancestry group. (**A**) Schematic of the rescue mechanisms where the modifying variant, a stop gained p.Tyr136Ter *in cis* with the dominant disease-causing p.Gly45Glu, results in incomplete penetrance by mono-allelic expression of the reference allele only (green). (**B**) Paired individual-level read data of the two variants occurring *in cis.* (**C**) East Asian (n=34) and Admixed American (n=1) individuals carrying both the pathogenic p.Gly45Glu variant (left panel) and the p.Tyr136Ter modifying variant (right panel).

### High rate of rescue identified for presumed disease-causing pLoF variants in genes associated with dominant disease

We sought to assess pLoF variants in genes associated with disease under the hypothesis that modifying variants could account for some observations of incomplete penetrance (Fig. 3a-b). We selected 77 haploinsufficient genes associated with severe, highly penetrant, early onset disorders (before the age of three years), expected to be absent or rarely seen in common disease studies or biobanks (e.g., gnomAD). We used *de novo* rate as a proxy for penetrance (Table S4). In these 77 genes, investigating 807,162 individuals, we found 4,464 pLoF variants, of which 3,957 were high-confidence pLoF variants by Loss-Of-Function Transcript Effect Estimator (LOFTEE)^4^, 3,223 in exomes (81%), 734 in genomes (19%, Fig. S3). The majority (87%), of these variants are extremely rare with an AC of five or less (Fig. S4).

We performed deep case-by-case assessments on the 734 high-confidence pLoF variants found in gnomAD genomes, building on a framework for pLoF variant assessment^25^ using conservative rules (Table S5) to exclude variants likely to not result in LoF (Table S6). Each pLoF variant was assigned a verdict of “Not LoF/Likely not LoF” when we found evidence the variant would not result in ablated expression of that protein (evade LoF), “Uncertain LoF” when there was partial or conflicting evidence or lack of data to assess the variant, and “LoF/Likely LoF” when we did not identify the reason for evading pLoF using this protocol. We defined 511 of 734 (69.6%) pLoF variants as explained (Not LoF/Likely not LoF; Fig. 3, blue). The most common explanations were: location in the last exon resulting in a variant not predicted to undergo nonsense-mediated decay (may or may not be pathogenic but excluded from this analysis); location in a region with low per-base expression score (pext, an average expression score derived from GTEx adult post-mortem tissues); rescue by modifying variants (multi-nucleotide variants or frame-restoring indels), location in a transcript containing a stop-codon (unlikely to be coding); and rescue by different types of splice modifying variants predicted by SpliceAI (in-frame up- or down-stream alternative splice site, no loss detected, or skipping/deletion of an in-frame exon; Fig. 3d). We labeled 86 of 734 (11.7%) pLoF variants as uncertain LoF (Fig. 3, grey), due to lack of read data limiting the ability to analyze for local modifying variants, concern for sequencing errors, contradictory SpliceAI predictions, or other types of weaker evidence of not being LoF (Fig. S5, Table S5). In 137 of 734 (18.7%) pLoF variants, the reason for lack of disease manifestation in 236 individuals was still unexplained and they remained interesting candidates for further investigation (Fig. 3, red, Table S6)

**Fig. 3:**
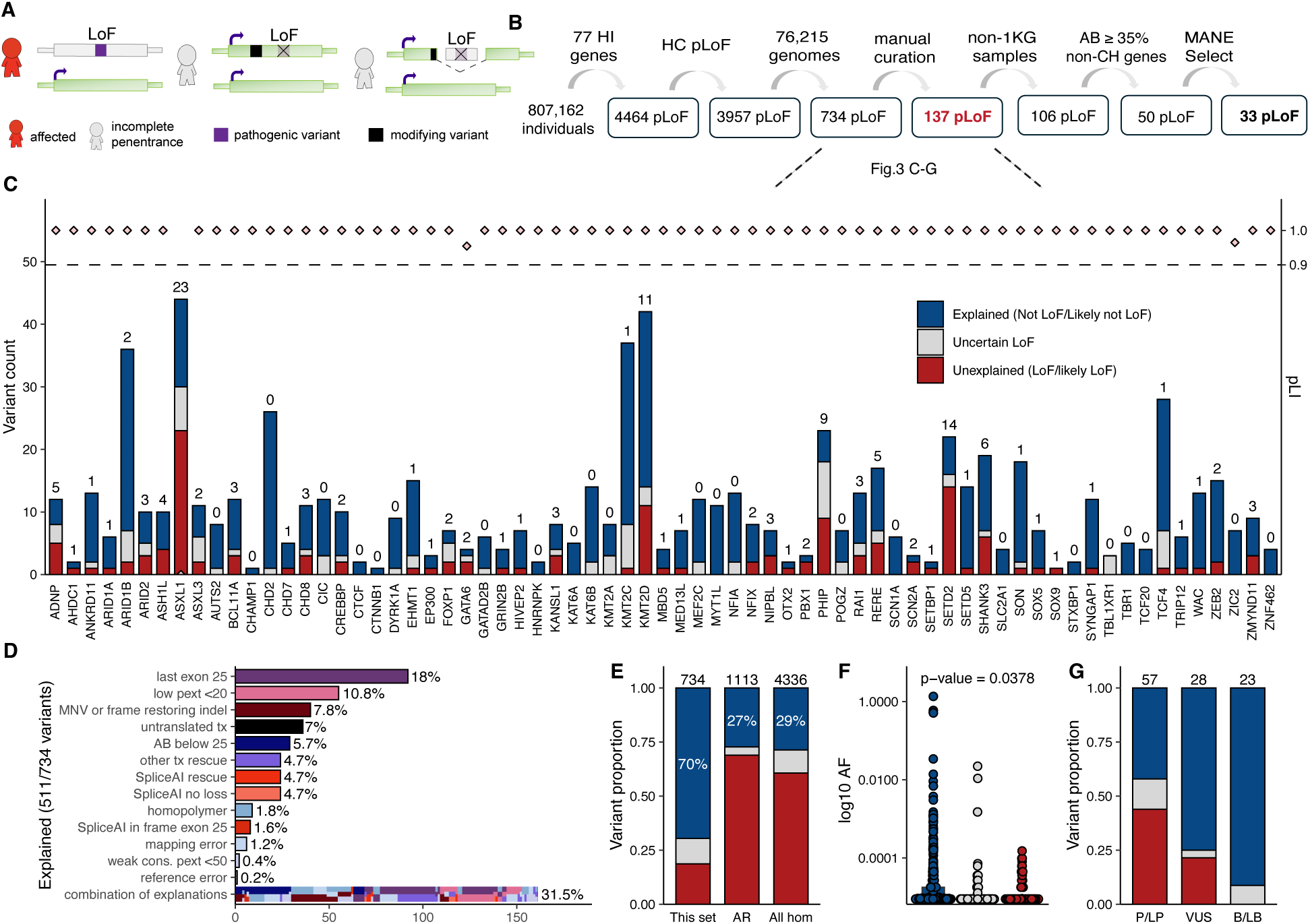
Deep assessment of 734 predicted high-confidence (HC) loss-of-function (pLoF) variants found in 77 haploinsufficient (HI) genes in 76,215 genomes. (**A**) Schematic of how modifying variants (black) may result in lack of a phenotype. (**B**) Filtering approach. 1KG: 1000 Genomes Project, AB: Allele balance, CH: clonal hematopoesis, MANE: Matched Annotation from NCBI and EMBL-EBI. (**C**) Variant count per HI gene colored by (also for E-G): explained (blue), uncertain (grey), unexplained (red). The number indicates unexplained variants (red). (**D**) The explanation for evading LoF in 511 of 734 (69.6%) of variants, MNV: multi-nucleotide variant, pext: per-base-expression score, tx: transcript, cons: conservation. (**E**) Comparison of outcome in different gene sets, this set (left), heterozygous pLoF variants in autosomal recessive (AR) disease genes in gnomAD v2 (middle), all homozygous (hom) pLoF variants in gnomAD v2 (right). (**F**) Allele frequency (AF) of variants in this set by LoF curation outcome (log10 scale). (**G**) The proportion of pLoF variants explained, uncertain, or unexplained within each ClinVar clinical classification category pathogenic/likely pathogenic (P/LP), uncertain significance (VUS), benign/likely benign (B/LB).

The observed pLoF evasion rate of 69.8% for this set of pLoF variants is notably higher than other sets of pLoF variants assessed in previous work reporting 27.3% (304/1,113) evasion for heterozygous pLoF variants associated with 22 AR disorders^25^, and 28.7% (1,245/4,336) in a set including all homozygous pLoF variants in gnomAD v2^4^ (Fig. 3e). The AF was higher for variants where LoF evasion could be explained compared to variants where the reason was yet to be found (p=0.0378, two-sided Student’s t-test; Fig. 3f).

We assessed how well the explained (blue) versus unexplained (red) variants align with reported ClinVar pathogenicity classifications. ClinVar classifications were available for 122 of 734 variants; B/LB variants were more likely to be explained compared to P/LP variants. The vast majority (21 of 23) of B/LB variants’ reason for evasion was explained by the protocol, and the remaining 2 of 23 were uncertain and excluded from further analysis. Of the 57/122 ClinVar pLoF variants that were P/LP, we could explain the lack of phenotype in gnomAD individuals for 25/57 variants (43.9%) (Fig. 3g, Table S7). Of all variants explained by being located in the last exon, located in low pext regions, or part of an MNVs in the gnomAD individuals, most were B/LB, with less than 25% of these variants being P/LP in ClinVar. In contrast, variants explained in our individuals due to potentially being sequencing errors (variant in homopolymer region) or somatic origin (allele balance of the alternate allele below 25%), are mostly reported as P/LP (Fig. S6), consistent with the expectation that these would result in LoF when they are bonafide germline variants.

After assessment using the LoF classification framework^25^, 137 variants in 236 individuals remained without an explanation. We noted a project-specific enrichment with 25.5% of variants (n=35) or 17.4% of samples (n=41), originating from the 1000 Genomes Project, although the 1000 Genomes Project constitutes less than 5% of gnomAD genome samples. Individuals from this cohort were thus excluded due to a concern that these may be cell culture-acquired variants (see discussion). For the remaining 106 variants in 195 individuals, we used skewed age to investigate signs of somatic origin which can rise to a high AF due to clonal hematopoiesis^29^. We observed a slightly higher age distribution in the 195 individuals compared to gnomAD genomes in general (Fig. S7a-b), and an even stronger skew for variants in clonal hematopoiesis-associated genes (n=36^29^; Fig. S7c). These variants were noted in the LoF classification framework to have a low AB, defined as an alternative allele ratio under 35% (with those under 20% already filtered by gnomAD QC practices and below 25% filtered by pLoF curation protocol) (Fig. S7d). After filtering variants found in clonal hematopoiesis genes and/or with low alternative AB due to possible somatic occurrence, only 50 variants in 104 individuals remained. Of these 50, 17 pLoF variants were not present in Matched Annotation from NCBI and EMBL-EBI (MANE) Select transcripts, suggesting that they might be of less biological relevance, leaving 33 of 734 (4.5%) pLoF variants as germline pLoF variation (Fig. S8).

### Incomplete penetrance due to alternative splicing mediated by non-coding sQTL variants

We investigated if alternative splicing of the region with a disease-causing pLoF, mediated by a specific sQTL, could explain incomplete penetrance in any of the original HC 734 pLoF variants identified in gnomAD (Fig. 4a). Out of the 734, nine pLoF variants (already labeled using the pLoF curation framework) fell in regions predicted to be alternatively spliced by a specific sQTL variant (Fig. 4b, Table S8). Notably, seven of the nine variants are found in regions with reduced pext scores, also suggesting alternative splicing in the general population, highlighting the usefulness of pext scores in assessing alternative splicing. Two *MEF2C* variants were marked as uncertain due to a 50% reduction in pext score and found in four genome-sequenced individuals with European ancestry, 5-88761023-C-A (n=3) and 5-88761110-C-A (n=1). Further investigation of these four individuals confirmed heterozygosity for the sQTL variant 5-89714113-C-G (non-coding) where the alternate allele is predicted to result in exon exclusion in three of four individuals (global AF 0.28, European AF 0.36), yet data for variant phasing was not available (Table S9). Investigation of *MEF2C* in all 807,162 individuals (including exomes) in gnomAD v4 displayed clustering of twelve pLoF variants in this region in combination with seven pLoF reported in ClinVar (four VUS and three P/LP), suggesting possible inaccurate P/LP classifications of the three ClinVar variants or incomplete penetrance of LoF variants in this region in some individuals due to alternative splicing (Fig. 4c, black box). Additionally, variants in *ARID1A* (n=3) and *ZEB2* (n=2) fell in a region with a ∼80% reduction in pext score (data not shown).

**Fig. 4:**
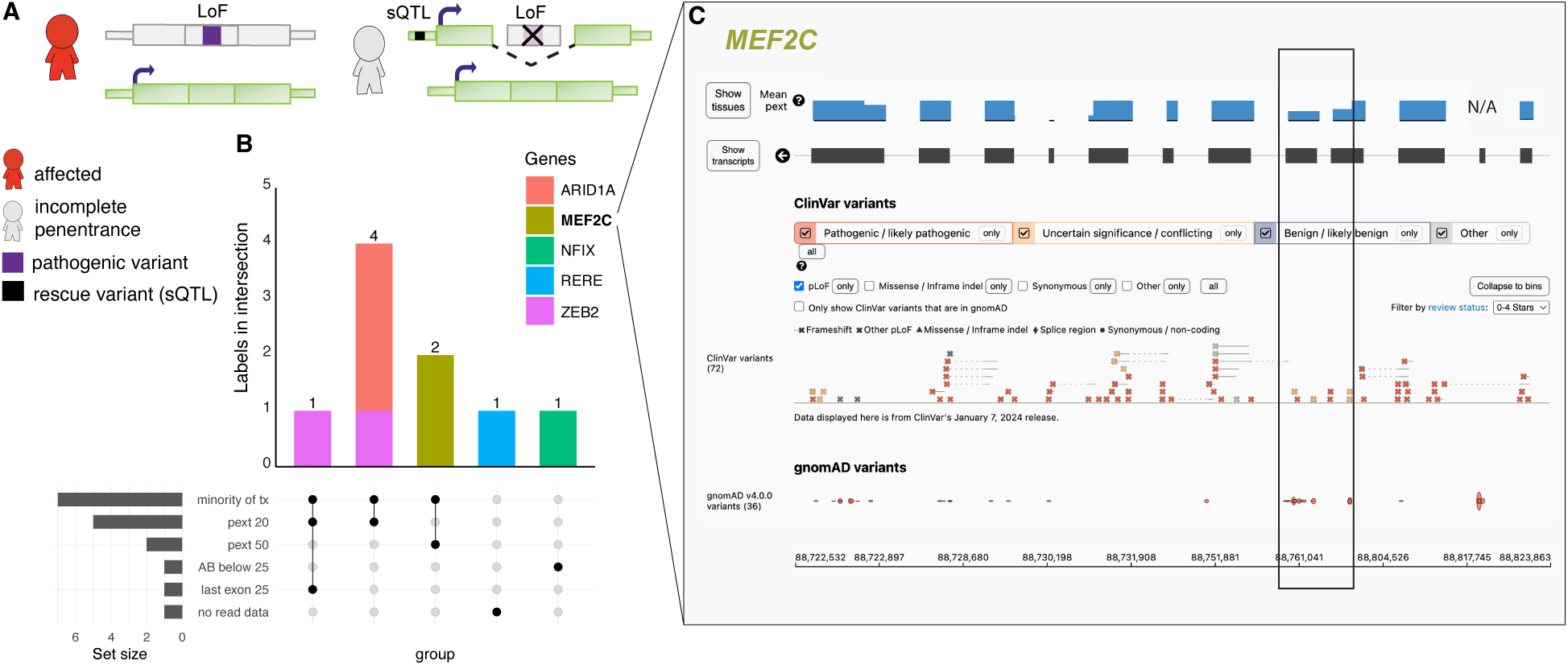
Splicing quantitative trait loci (sQTL) analysis. (**A**) Schematic of alternative splicing as a mechanism for incomplete penetrance. (**B**) Labels for each of the nine predicted loss of function (pLoF) variants. Bars are colored by variant count per gene. Tx: transcript, pext: per base expression score, AB: allele balance. (**C**) Overview of pLoF variants in *MEF2C* in ClinVar and in gnomAD v4 (all variants, including exomes). Black box marks region of reduced mean pext score (from v2, blue bar, for one exon the score is not available [N/A]) and enrichment of *MEF2C* pLoF variants in gnomAD.

## DISCUSSION

Presumably unaffected individuals with variants predicted to be associated with severe, highly penetrant, and early-onset disease present an opportunity to improve understanding of variant pathogenicity, interpretation, and penetrance. Previous studies of disease-variants present in population databases have focused on descriptive reports^1^, statistical associations between disease-variants and phenotypes^8–10,14,30^, estimated penetrance of specific disorders^8,9,31^, gene-specific mechanisms^15^ or a specific type of modifying variants (e.g. eQTLs/ sQTLs)^16,18,32^. We aimed to move beyond previous efforts by using a large-scale but non-statistical approach, under the hypothesis that some of these modes are rare and need in-depth investigation of the relevant region to allow discovery of the underlying reason for the lack of disease manifestation. We investigated individuals on a case-by-case level and identified a wide range of explanations underlying tolerance for these pLoF variants in a large set of distinct disorders.

Using the gnomAD v4 dataset with variant data from 807,162 individuals, we could confirm that large-scale efforts like gnomAD increasingly include rare clinically relevant variants, underlining that continued aggregation of data, especially focusing on diversity and underrepresented groups, will power variant analysis. Increased sample size provides more accurate predictions of variant rarity, as well as improved coverage of less common variation. Allele frequencies are mostly consistent over gnomAD versions and allele counts for P/LP ClinVar variants and pLoF variants in genes associated with severe disease are consistently low. The more than 5-fold increase in individuals between v2 and v4 does result in more unique P/LP variants as well as higher allele counts in some cases, which is expected with the larger database size and needs to be considered in analysis.

The utility of studying individuals with pathogenic variants that do not present with the associated phenotype, and the importance of diverse ancestry representation in any population database, was demonstrated by a case of genetic ancestry group-specific incomplete penetrance in East Asians of a *GJB2* variant associated with a lethal form of KID syndrome. We note that although bi-allelic loss of *GJB2* is associated with AR hearing loss, there is a high observed/expected number of pLoF variants in *GJB2* in gnomAD, with a LOEUF score of 1.97, higher than 99.9% (5/3064) of AD and/or AR genes, suggesting a possible selective advantage of pLoF variants in *GJB2* which has been explored previously^33–35^.

We investigated all pLoF found in 76,215 v4 genomes in 77 genes associated with autosomal dominant, severe, highly penetrant, early-onset disease. As expected, 76 of 77 genes were loss-of-function constrained with a pLI ≥0.9 in gnomAD v4. *ASXL1* has a pLI of 0.0 in both v4 and v2 due to somatic variants rising to higher allele balance due to clonal hematopoiesis and deflating the pLI constraint score^2^. Of 734 pLoF variants identified in the 77 genes, we found an explanation for the lack of disease manifestation and tolerance of the pLoF variant for 95% of variants. The explanations identified highlighted a combination of our current limitations in pLoF annotations, the occurrence of somatic variation, rare instances of sequencing artifacts, and examples of incomplete penetrance of pathogenic variants. The most common explanations were location in the last exon (or last 50 base pairs of the penultimate exon), low pext region, rescue by nearby secondary variants (MNVs/frame restoring indels or splice variants), and somatic variants resembling germline variants due to clonal expansion resulting in elevated allele balance of the alternate allele. In general, a pLoF variant in this set of genes associated with highly penetrant conditions can be explained by careful evaluation described here, and unexplained incomplete penetrance is very rare.

The first line of pLoF assessment in this study was based on our framework for loss-of-function curation^21^, which evaluates if there is reason to believe the pLoF variants do not result in ablated protein expression. The verdict “Not LoF/Likely not LoF” does not determine variant pathogenicity, e.g., a 3’ pLoF variant not resulting in lost protein expression can still be pathogenic by expression of a truncated protein. Of the 734 pLoF variants evaluated with this framework, 122 were also reported in ClinVar and allowed us to demonstrate that indeed the majority of explained variants in the last exon, low pext regions, or part of multinucleotide variants do not result in LoF and are benign, and only under rare circumstances P/LP. Pathogenicity must be assessed considering specific gene properties, such as the presence of a functional domain or nearby established pathogenic variants. Further, a lowly expressed region (low pext score region) that shows evidence of biological relevance, e.g., expression in a disease-relevant tissue, can still have important implications in disease. The current restriction to adult tissues in GTEx, and lack of any pediatric or prenatal tissues, is limiting when interpreting transcript properties and expression of genes causing syndromes manifesting prenatally.

Somatic occurrence and clonal expansion over time likely explain some germline-resembling unexpected variants. For one, we observed an overrepresentation of pLoF variants in severe pediatric disease genes in samples from the 1000 Genomes project, and to a lesser extent, a general enrichment for (germline blood) samples originating from cancer cohorts.

Samples from the 1000 Genomes project are derived from cultured cell lines, suggesting that these variants may be somatic variants that have gone through clonal expansion over time (aging), rather than germline in origin. This highlights the need for caution when using non-primary cells to study the landscape of human variation tolerance and is one of the reasons that the 1000 Genomes Project (1KG) allele information is available on each variant page as a separate tab of the gnomAD v4 frequencies table, allowing identification of variants originating from this project. Second, we observed skewed age distribution towards elderly individuals of samples with variants in clonal hematopoiesis genes, as well as variants labeled because of low alternative allele balance, demonstrating how skewed age can help guide the identification of somatic occurrence of variants, as shown previously^29^.

We found that 9 of 734 pLoF variants fell in exons that have been associated with alternative splicing mediated by specific sQTL variants. sQTLs have previously been suggested to play a role in penetrance. Einson et al., showed that natural selection acts on haplotype configurations that reduce the transcript inclusion of putatively pathogenic variants in TOPMed consortium data, especially when limiting to haploinsufficient genes^17^, and Beaumont et al., reported a general observation of the non-uniform distribution of pLoF variants in haploinsufficiency disease-genes and suggested alternative splicing and translation re-initiation of these regions could explain incomplete penetrance^36^. We present a specific example of an sQTL variant associated with alternative splicing of a certain region of *MEF2C. MEF2C* haploinsufficiency is associated with a neurodevelopmental disorder presenting with developmental and cognitive delay, limited language and walking, hypotonia, and seizures (MIM: 600662)^37^. The combination of clustering of pLoF variants gnomAD (twelve pLoF) and ClinVar P/LP variants (three P/LP and four VUS pLoF) could suggest that pLoF in this region can be of incomplete penetrance in the gnomAD individuals, potentially by alternative splicing that could be mediated by the sQTL carried by three out of four individuals with sQTL genotype data and pLoF variants in this region. It is also possible that the alternative splicing is population-wide, pLoF in this region is not associated with disease, and the P/LP variants in this region are examples of misclassifications in ClinVar.

This study helps further our understanding and ability to interpret variant effects and understand incomplete penetrance of disease, especially focusing on pLoF variants. Most importantly, we highlight the complicated assessment of pLoF variants effect, which is a major challenge in variant classification, interpretation, diagnostic testing, and genetic risk prediction. Most presumed disease-causing pLoF variants here and in other population databases can be reclassified as misannotations, somatic, or artifacts by deep investigation, but also include cases of incomplete penetrance mediated by other modifying genetic variants. Only 4.5% of pLoF variants associated with the 77 high-penetrant conditions were unexplained, highlighting that deep assessment on a variant-by-variant basis far beyond standard high-throughput pipelines is useful and needed. Failing to do so runs the risk of overinterpreting pathogenicity in clinic and in research, especially for severe conditions in unaffected individuals where the false discovery rate is higher^38^.

Although not within the scope of this study, molecular assessment of these unexpected cases will be useful next steps and allow a deeper understanding of biological mechanisms underlying disease-penetrance. Continued aggregation of sequencing data, ideally associated with phenotype, will allow further studies in this area. Especially when focusing on under-represented groups where yet undiscovered haplotypes with modifying events, combined with known disease variants, can inform new mechanisms resulting in incomplete penetrance of disease.

## ONLINE METHODS

### ClinVar variants in gnomAD

ClinVar variants reported here include all variants available for download by December 1st, 2023 (https://ftp.ncbi.nlm.nih.gov/pub/clinvar/tab_delimited/) filtered to indels less than 50 base pairs and single nucleotide variants on chromosomes 1:22+X+Y. All variants were grouped based on the aggregate ClinVar classification category (clinical significance) into B/LB when reported as “Likely benign”, “Benign” or “Benign/Likely benign”, Uncertain when “Uncertain significance”, P/LP when “Pathogenic”, “Likely pathogenic” or “Pathogenic/Likely pathogenic” and conflicting when “Conflicting interpretations of pathogenicity”. A small proportion of variants that did not fall into any of the above-listed classification categories were excluded; in the majority of these the clinical significance had not been provided (“not provided”/ “no interpretation for the single variant”) followed by “drug response”, “risk factor” and “association”. For “ClinVar variants present in gnomAD” we included any ClinVar variant represented in the 807,162 individuals that passed gnomAD QC filters^20^, only including a variant if “pass” in either exomes or genomes. A combined AF was calculated for variants detected with both exome and genome sequencing. For variants only identified by one sequencing method the AF of sample sequenced by that method was used. The inheritance pattern for each pathogenic variant was determined using the reported inheritance in OMIM of the relevant gene (by October 23, 2023) categorized as AR, AR/AD (if both patterns were reported), AD or XL. Variants that fell in genes not reported in OMIM, genes with no reported inheritance, or other types/combinations of inheritance were excluded from the inheritance analysis (Fig. 1f).

### Rescue by local pLoF events in a subset of P/LP in ClinVar

We investigated if a set of P/LP variants were tolerated due to modifying pLoF variants in the same gene, potentially reducing the expression of the pathogenic allele. We included all P/LP ClinVar variants found in 807,162 individuals in gnomAD that were located in a gene not constrained for LoF (predicted loss-of-function intolerance <0.9, pLI) and with at least one condition of AD inheritance in OMIM, suggesting that pathogenicity is more likely to act through a gain-of-function mechanism, found in 50 individuals or fewer. The pLI score is mostly stable over v2 and v4 versions and its dichotomous nature makes it suitable for determining haploinsufficiency in the context of variant depletion in a population database. Of note, the Loss-of-function Observed/Expected Upper-bound Fraction (LOEUF) score, a continuous metric of pLoF depletion, shows some variance between versions due to sample size increases, resulting in an increased discovery rate of ultra-rare variants along with an increased number of artifacts (Fig. S9). We then manually assessed all pairs of P/LP variants occurring in combination with a pLoF event with a global AF equal or less than 1% (AF ≤ 0.01) that passed gnomAD QC filters (depth < 10, genotype quality < 20, minor allele balance < 0.2 for alternate alleles of heterozygous genotypes). Combinations were determined interesting if passing three criteria (1) carriership of a pLoF variant in all individuals with the P/LP variant (2) a pLoF variant determined as a true LoF resulting in ablated protein product using LoF curation explained below (filtering artifacts, somatic variants, missanotations or rescued variant), (3) a P/LP variant acting through dominant gain-of-function mechanism.

### Severe disease genes investigated in gnomAD

We assessed over 450 genes associated with rare disorders starting from gene lists from resources like the Deciphering Developmental Disorders (DDD) study and the ClinGen Dosage sensitivity curation project as well as internally collected genes and genes shared by collaborators. By literature review, we filtered the >450 genes to a stringent gene-disease list where all genes were associated with disorders that met the following criteria: (1) Caused by autosomal haploinsufficiency in at least three unrelated cases. (2) Early-onset, defined as before the age of three. (3) Severe phenotype unlikely to be compatible with participation in common disease studies or prohibit consent to such study, i.e., mainly inclusion of syndromes resulting in severe congenital malformations and/or neurodevelopmental symptoms of severe degree. (4) Highly penetrant and limited variable expressivity of phenotypes, scored as estimated penetrance >70%, >80%, >90% or 100%. For many disorders there is no clear penetrance estimate available in literature or within resources like GeneReviews. For these disorders, the reported proportion of de novo versus inherited variants was used as a proxy for penetrance.

### Loss-of-function curation and splicing quantitative trait loci analysis

We included all pLoF variants (nonsense, frameshift, essential splice variants) found in v4.1.0 genomes (76,215 individuals) in any GENCODE 39 transcripts that were high-confidence according to the Loss-Of-Function Transcript Effect Estimator^4^. LoF variants were assessed using the framework for LoF curation previously published by this group^25^, adapted for this project. In general, we used conservative thresholds to exclude any pLoF variants likely not to cause loss of function. The specific set of rules used for this project are found in Table S5. Splice variants were assessed for rescues (within 1000 bp) using SpliceAI lookup (https://spliceailookup.broadinstitute.org/). pLoF variants labeled as Not LoF or Likely not LoF were grouped and categorized as “Explained” (blue), variants labeled as Uncertain were excluded (grey), and variants scored as LoF or Likely LoF were grouped and referred to as “unexplained” (red).

Splicing Quantitative Trait Loci (sQTL) analysis was performed on pLoF variants in v4 genomes according to methods previously described^17^. In short, cis-sQTL variants were identified from GTEx v8 data by association between exon inclusion levels (PSI) and genetic variants in 1Mb window, at <5% FDR.

### Statistical analysis and data visualization

All statistical analysis and data visualization for figures were generated using R v4.3.1 (https://www.r-project.org/), mainly utilizing libraries from ggplot2 v3.4.3, ComplexUpset v1.3.3, and ggpubr v0.6.0.

## DATA AVAILABILITY

Variant data, constraint metrics for gnomAD v4 and v2, and loss of function curation data are publicly available at https://gnomad.broadinstitute.org/downloads.

## ACKNOWLEDGMENTS

We thank the individuals and researchers whose data is in gnomAD for their contributions. We thank Dr. Mark Daly, Sinéad Chapman, Caroline Cusick, and the team for sharing individual-level sequencing read data for the *GJB2* case. S.G. was supported by The Knut and Alice Wallenberg Foundation scholarship program for postdoctoral studies at the Broad Institute. SLS was supported by a fellowship from the Manton Center for Orphan Disease Research at Boston Children’s Hospital. DGM is supported by a National Health and Medical Research Council (Australia) investigator grant 2009982. The study was funded in part by The Mather’s Foundation to SG and AODL and by the NIH/NHGRI grant U24HG011450 to HLR.

## AUTHOR CONTRIBUTIONS

SG planned and conducted all analyses, wrote the manuscript, and prepared all figures and tables. MS-B and NAW contributed to LoF curation. SLS, JKG, and MW contributed to analyses on gnomAD data. JE and TL performed the sQTL analysis. HR, DGM, AOD-L contributed with study design, funding, and feedback. All authors approved the final version of the manuscript.

## REFERENCES

1. Tarailo-Graovac, M., Zhu, J. Y. A., Matthews, A., van Karnebeek, C. D. M. & Wasserman, W. W. Assessment of the ExAC data set for the presence of individuals with pathogenic genotypes implicated in severe Mendelian pediatric disorders. Genet. Med. 19, 1300–1308 (2017).

2. Carlston, C. M. et al. Pathogenic ASXL1 somatic variants in reference databases complicate germline variant interpretation for Bohring-Opitz Syndrome. Hum. Mutat. 38, 517–523 (2017).

3. Lek, M. et al. Analysis of protein-coding genetic variation in 60,706 humans. Nature 536, 285–291 (2016).

4. Karczewski, K. J. et al. The mutational constraint spectrum quantified from variation in 141,456 humans. Nature 581, 434–443 (2020).

5. Wright, C. F. et al. Assessing the Pathogenicity, Penetrance, and Expressivity of Putative Disease-Causing Variants in a Population Setting. Am. J. Hum. Genet. 104, 275–286 (2019).

6. Sorscher, S. Ascertainment Bias and Estimating Penetrance. JAMA Oncol 4, 587 (2018).

7. Kingdom, R. & Wright, C. F. Incomplete Penetrance and Variable Expressivity: From Clinical Studies to Population Cohorts. Front. Genet. 13, 920390 (2022).

8. Minikel, E. V. et al. Quantifying prion disease penetrance using large population control cohorts. Sci. Transl. Med. 8, 322ra9 (2016).

9. Goodrich, J. K. et al. Determinants of penetrance and variable expressivity in monogenic metabolic conditions across 77,184 exomes. Nat. Commun. 12, 3505 (2021).

10. Patel, K. et al. Penetrance of MODY is substantially lower in clinically unselected cohort: important implications for opportunistic genomic testing. in DIABETOLOGIA vol. 65 S41– S41 (SPRINGER ONE NEW YORK PLAZA, SUITE 4600, NEW YORK, NY, UNITED STATES, 2022).

11. Mao, K. et al. FOXI3 pathogenic variants cause one form of craniofacial microsomia. Nat. Commun. 14, 2026 (2023).

12. Michaud, V., et al. The contribution of common regulatory and protein-coding TYR variants in the genetic architecture of albinism. bioRxiv (2021) doi:10.1101/2021.11.01.21265733.

13. Timberlake, A. T. et al. Two locus inheritance of non-syndromic midline craniosynostosis via rare SMAD6 and common BMP2 alleles. Elife 5, (2016).

14. Tuke, M. A. et al. Mosaic Turner syndrome shows reduced penetrance in an adult population study. Genet. Med. 21, 877–886 (2019).

15. Naqvi, S. et al. Precise modulation of transcription factor levels identifies features underlying dosage sensitivity. Nat. Genet. (2023) doi:10.1038/s41588-023-01366-2.

16. Castel, S. E. et al. Modified penetrance of coding variants by cis-regulatory variation contributes to disease risk. Nat. Genet. 50, 1327–1334 (2018).

17. Einson, J. et al. Genetic control of mRNA splicing as a potential mechanism for incomplete penetrance of rare coding variants. Genetics 224, (2023).

18. Wigdor, E. M. et al. Investigating the role of common*cis*-regulatory variants in modifying penetrance of putatively damaging, inherited variants in severe neurodevelopmental disorders. medRxiv (2023) doi:10.1101/2023.04.20.23288860.

19. Bamshad, M. J., Nickerson, D. A. & Chong, J. X. Mendelian Gene Discovery: Fast and Furious with No End in Sight. Am. J. Hum. Genet. 105, 448–455 (2019).

20. Gudmundsson, S. et al. Variant interpretation using population databases: Lessons from gnomAD. Hum. Mutat. 43, 1012–1030 (2022).

21. Singer-Berk, M. et al. Advanced variant classification framework reduces the false positive rate of predicted loss of function (pLoF) variants in population sequencing data. medRxiv (2023) doi:10.1101/2023.03.08.23286955.

22. Landrum, M. J. et al. ClinVar: improvements to accessing data. Nucleic Acids Res. 48, D835–D844 (2020).

23. Richards, S. et al. Standards and guidelines for the interpretation of sequence variants: a joint consensus recommendation of the American College of Medical Genetics and Genomics and the Association for Molecular Pathology. Genet. Med. 17, 405–424 (2015).

24. ClinVar website. ClinVar submission statistics. https://www.ncbi.nlm.nih.gov/clinvar/submitters/.

25. Singer-Berk, M. et al. Advanced variant classification framework reduces the false positive rate of predicted loss-of-function variants in population sequencing data. Am. J. Hum. Genet. 110, 1496–1508 (2023).

26. Jonard, L. et al. A familial case of Keratitis-Ichthyosis-Deafness (KID) syndrome with the GJB2 mutation G45E. Eur. J. Med. Genet. 51, 35–43 (2008).

27. Ogawa, Y. et al. Revertant mutation releases confined lethal mutation, opening Pandora’s box: a novel genetic pathogenesis. PLoS Genet. 10, e1004276 (2014).

28. Sakata, A. et al. Hearing and Hearing Loss Progression in Patients with GJB2 Gene Mutations: A Long-Term Follow-Up. Int. J. Mol. Sci. 24, (2023).

29. Gudmundsson, S., Carlston, C. M. & O’Donnell-Luria, A. Interpreting variants in genes affected by clonal hematopoiesis in population data. Hum. Genet. (2023) doi:10.1007/s00439-023-02526-4.

30. DeBoever, C. et al. Medical relevance of protein-truncating variants across 337,205 individuals in the UK Biobank study. Nat. Commun. 9, 1612 (2018).

31. de Masfrand, S. et al. Penetrance, variable expressivity and monogenic neurodevelopmental disorders. Eur. J. Med. Genet. 104932 (2024).

32. Einson, J. et al. Genetic control of mRNA splicing as a potential mechanism for incomplete penetrance of rare coding variants. bioRxiv 2023.01.31.526505 (2023) doi:10.1101/2023.01.31.526505.

33. Common, J. E. A., Di, W.-L., Davies, D. & Kelsell, D. P. Further evidence for heterozygote advantage of GJB2 deafness mutations: a link with cell survival. J. Med. Genet. 41, 573– 575 (2004).

34. D’Adamo, P. et al. Does epidermal thickening explain GJB2 high carrier frequency and heterozygote advantage? Eur. J. Hum. Genet. 17, 284–286 (2009).

35. Vuckovic, D. et al. Connexin 26 variant carriers have a better gastrointestinal health: is this the heterozygote advantage? Eur. J. Hum. Genet. 23, 563 (2015).

36. Beaumont, R. N., Hawkes, G., Gunning, A. C. & Wright, C. F. Clustering of predicted loss-of-function variants in genes linked with monogenic disease can explain incomplete penetrance. Genome Med. 16, 64 (2024).

37. Cooley Coleman, J. A., et al. Clinical findings from the landmark MEF2C-related disorders natural history study. Mol Genet Genomic Med 10, e1919 (2022).

38. Gudmundsson, S. et al. Addendum: The mutational constraint spectrum quantified from variation in 141,456 humans. Nature (2021) doi:10.1038/s41586-021-03758-y.

39. Moreno-Pelayo, M. A. et al. A cysteine substitution in the zona pellucida domain of alpha-tectorin results in autosomal dominant, postlingual, progressive, mid frequency hearing loss in a Spanish family. J. Med. Genet. 38, E13 (2001).

