## Supplementary Information for "Exploring penetrance of clinically relevant variants in over 800,000 humans from the Genome Aggregation Database"

### Exploring disease-variants and variable penetrance in over 800,000 individuals from the Genome Aggregation Database

### Supplementary Notes

#### Note S1: Rescue by local pLoF events in a subset of P/LP in ClinVar

Of 3957 of P/LP variants (found in in 31,014 of 807,162 individuals in gnomAD), 90 unique P/LP variants in 222 individuals occurred in combination with a pLoF (AF ≤ 0.01, depth < 10, genotype quality < 20, minor allele balance < 0.2) in the same gene. Evaluation following three steps revealed one interesting case: a pLoF in combination with an otherwise lethal autosomal dominant variant in *GJB2* (see results). Additional to the highlighted GJB2 case (results), one P/LP variant (11-121166704-G-A) in *TECTA* associated with autosomal non-syndromic dominant hearing loss reportedly acting through dominant negative effect[39] was rescued by an MNV variant where the secondary 11-121166705-C-A variant occurring in the same codon as the P/LP variant altered the missense variant to a nonsense variant in one individual. Biallelic nonsense variants in *TECTA* are associated with autosomal recessive hearing loss, and thus this heterozygous MNV is not disease-causing in this one individual. We also found one combination of a P/LP variant (7-87453031-C-T) in *ABCB4* gene reported to cause Intrahepatic cholestasis of pregnancy in combination with a pLoF splice variant in 17 individuals of European ancestry. The other 87 unique P/LP variants occurring in combination with a pLoF were excluded based on (1) carriership of a pLoF variant was not found in the majority (≤50%) of individuals with the P/LP variant (79 variants), (2) the pLoF variant showed evidence of being "not/likely not pLoF" and thus not resulting in ablated protein product (6 variants), and (3) the P/LP variant was not acting through dominant gain-of-function mechanism (2 variant) (Table S3).

### Supplementary Figures

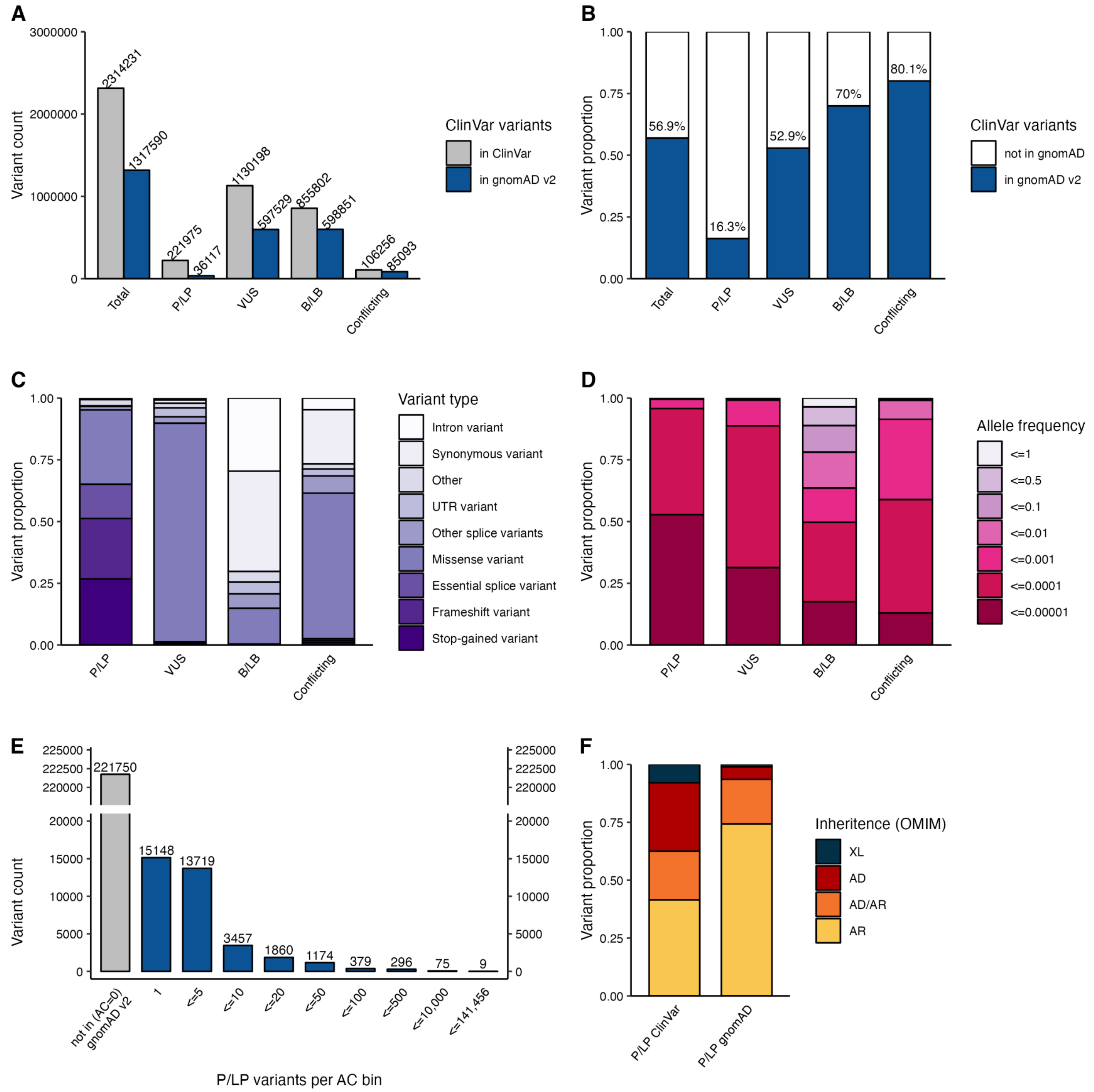


**Fig. S1:** Representation of ClinVar variants in 141,456 individuals in gnomAD v2. (A) Variant count of ClinVar variants in ClinVar (grey) vs. gnomAD (blue) in each classification category pathogenic/likely pathogenic (P/LP), variant of uncertain significance (VUS), benign/likely benign (B/LB) or with conflicting classifications. (B) Percentage of ClinVar variants reported in gnomAD in at least one individual. (C) The proportion of variants by variant type within each clinical significance classification and, (D) within each allele frequency bin. (E) Total number of P/LP variants within each allele count bin. (F) The inheritance pattern of the gene harboring the P/LP variants in gnomAD vs. inheritance of all variants in ClinVar.

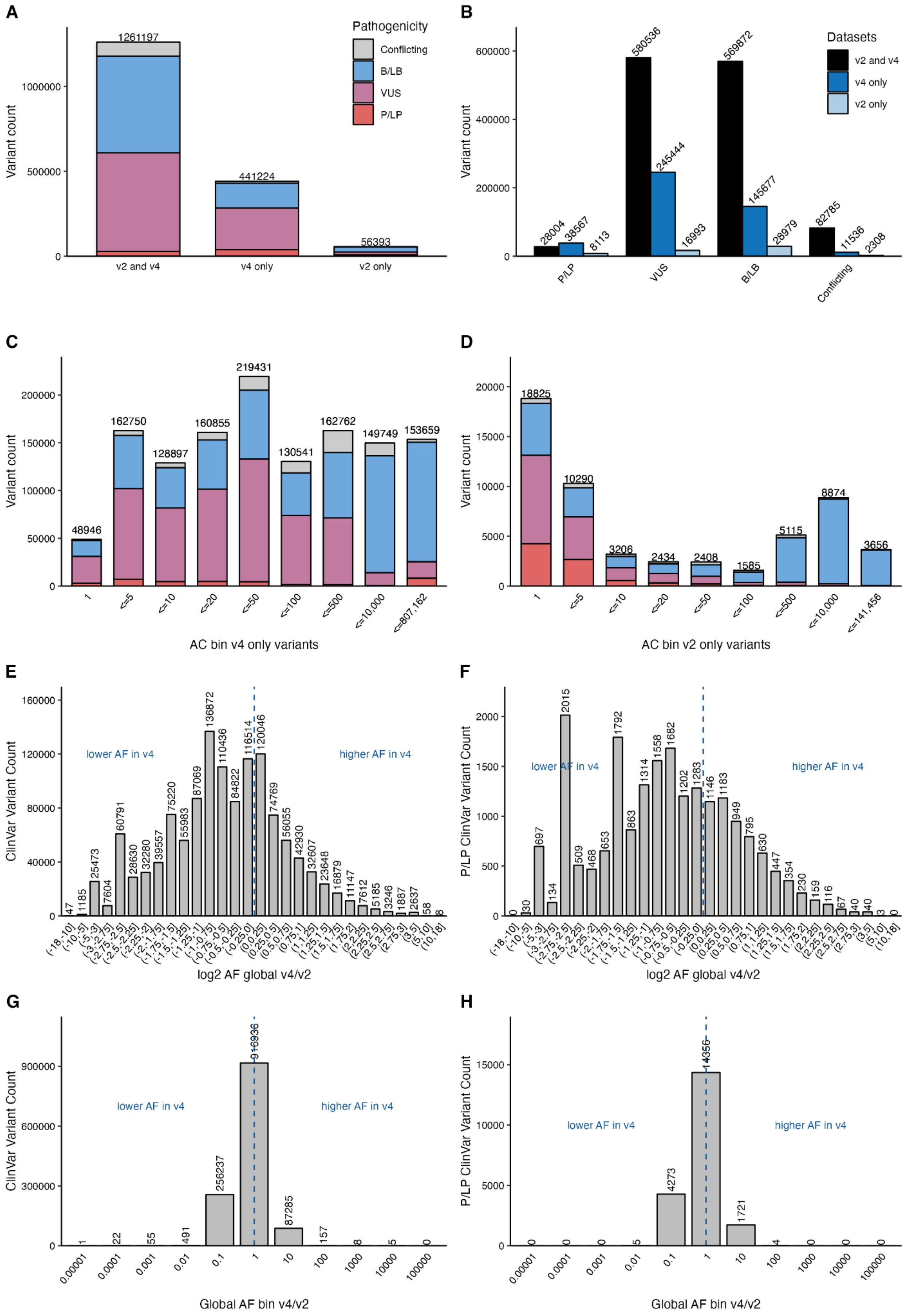
A
Variant count
1261197
441224
56393
v2 and v4
v4 only
v2 only
Pathogenicity
Conflicting
B/LB
VUS
P/LP
B
Variant count
Datasets
v2 and v4
v4 only
v2 only
28004
38567
8113
580536
245444
16993
569872
145677
28979
82785
11536
2308
P/LP
VUS
B/LB
Conflicting
C
Variant count
48946
162750
128897
160855
219431
130541
162762
149749
153659
1
<=5
<=10
<=20
<=50
<=100
<=500
<=10,000
<=807,162
AC bin v4 only variants
D
Variant count
18825
10290
3206
2434
2408
1585
5115
8874
3656
1
<=5
<=10
<=20
<=50
<=100
<=500
<=10,000
<=141,456
AC bin v2 only variants
E
ClinVar Variant Count
lower AF in v4
higher AF in v4
47
1185
25473
7604
60791
28630
32280
39557
75220
55983
87069
136872
110436
84822
116514
120046
74769
56055
42930
32607
23648
16879
11147
7612
5185
3246
1887
2637
58
8
log2 AF global v4/v2
F
P/LP ClinVar Variant Count
lower AF in v4
higher AF in v4
0
30
697
134
2015
509
468
653
1792
863
1314
1558
1682
1202
1283
1146
1183
949
795
630
447
354
230
159
116
67
40
40
3
0
log2 AF global v4/v2
G
ClinVar Variant Count
lower AF in v4
higher AF in v4
1
22
55
491
256237
916936
87285
157
8
5
0
0.00001
0.0001
0.001
0.01
0.1
1
10
100
1000
10000
100000
Global AF bin v4/v2
H
P/LP ClinVar Variant Count
lower AF in v4
higher AF in v4
0
0
0
5
4273
14356
1721
4
0
0
0
0.00001
0.0001
0.001
0.01
0.1
1
10
100
1000
10000
100000
Global AF bin v4/v2

**Fig. S2:** ClinVar variants represented in gnomAD v2 and v4. (A-B) Total ClinVar variant count per dataset grouped by pathogenicity. (C-D) Allele count (AC) bins for variants only present in v4 (C) or v2 (D). (E-F) Allele frequency (AF) ratio v4 compared to v2 (log2 scale) all ClinVar variants (E) and ClinVar P/LP variants (F) that are present in both v2 and v4 datasets. (G-H) The ratio between AF bins in v4 compared to v2 all ClinVar variants (G) and ClinVar P/LP variants (H) that are present in both v2 and v4 datasets. The majority of variants belong to the same AF bin in both datasets, while 21% (n=4278) of P/LP variants are 10 times less common (equal to or below 0.1) in v4.

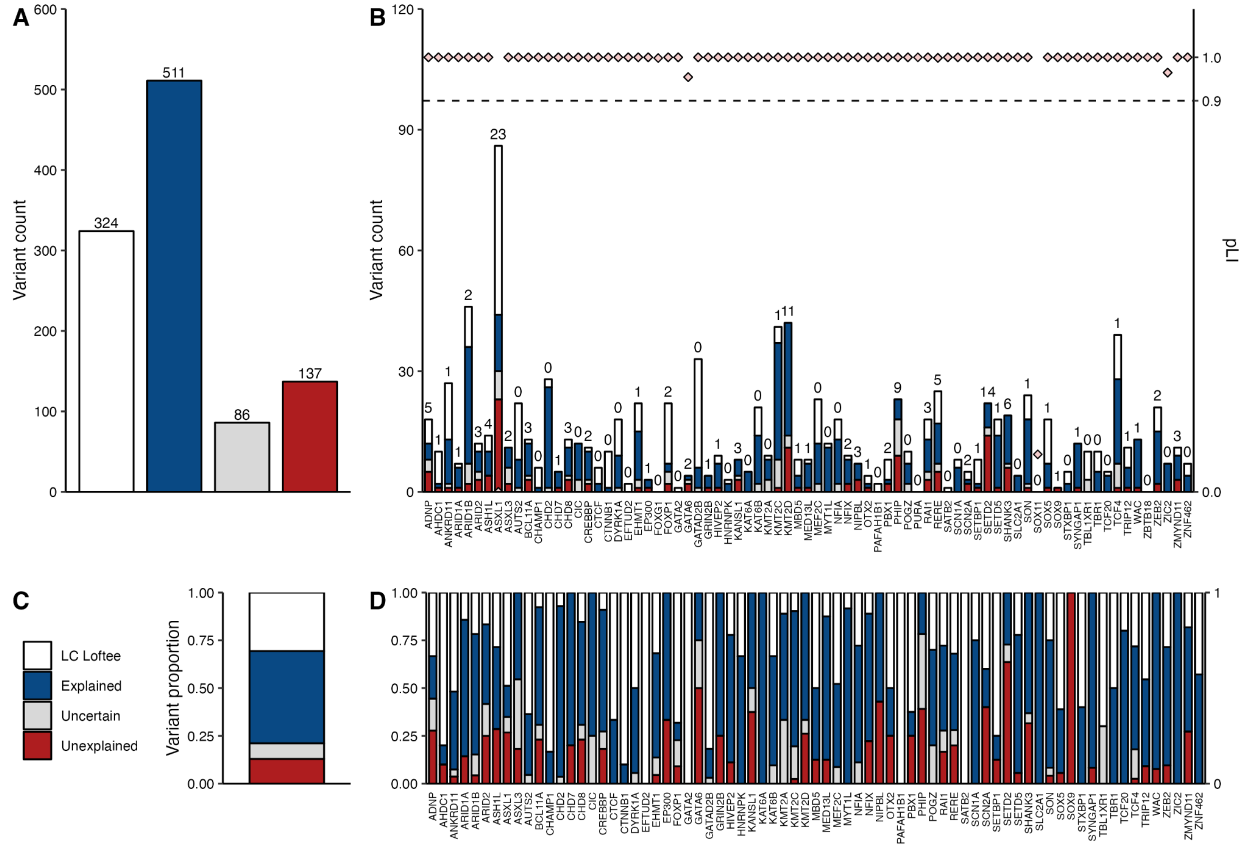


**Fig. S3:** Investigation of all pLoF variants (n=1,058) in 77 genes associated with severe, early-onset highly penetrance disease. Loss-Of-Function Transcript Effect Estimator (LOFTEE) filtered 30.6% (324/1,058) of pLoF variants as low-confidence (LC; white). Application of the Singer-Berk et al., pLoF framework led to 48.3% (511/1,058) assessed as likely not/not pLoF and thus considered explained (blue), and 8.1% (86/1,058) excluded as uncertain (grey). Of 1058 pLoF variants, 12,9% (n=137) the lack of disease manifestation in gnomAD individuals was still unexplained (red) and subject to further evaluation. (A) Variant counts per category across all variants, (B) by gene, and (C) all variants displayed as proportion (D) per gene.

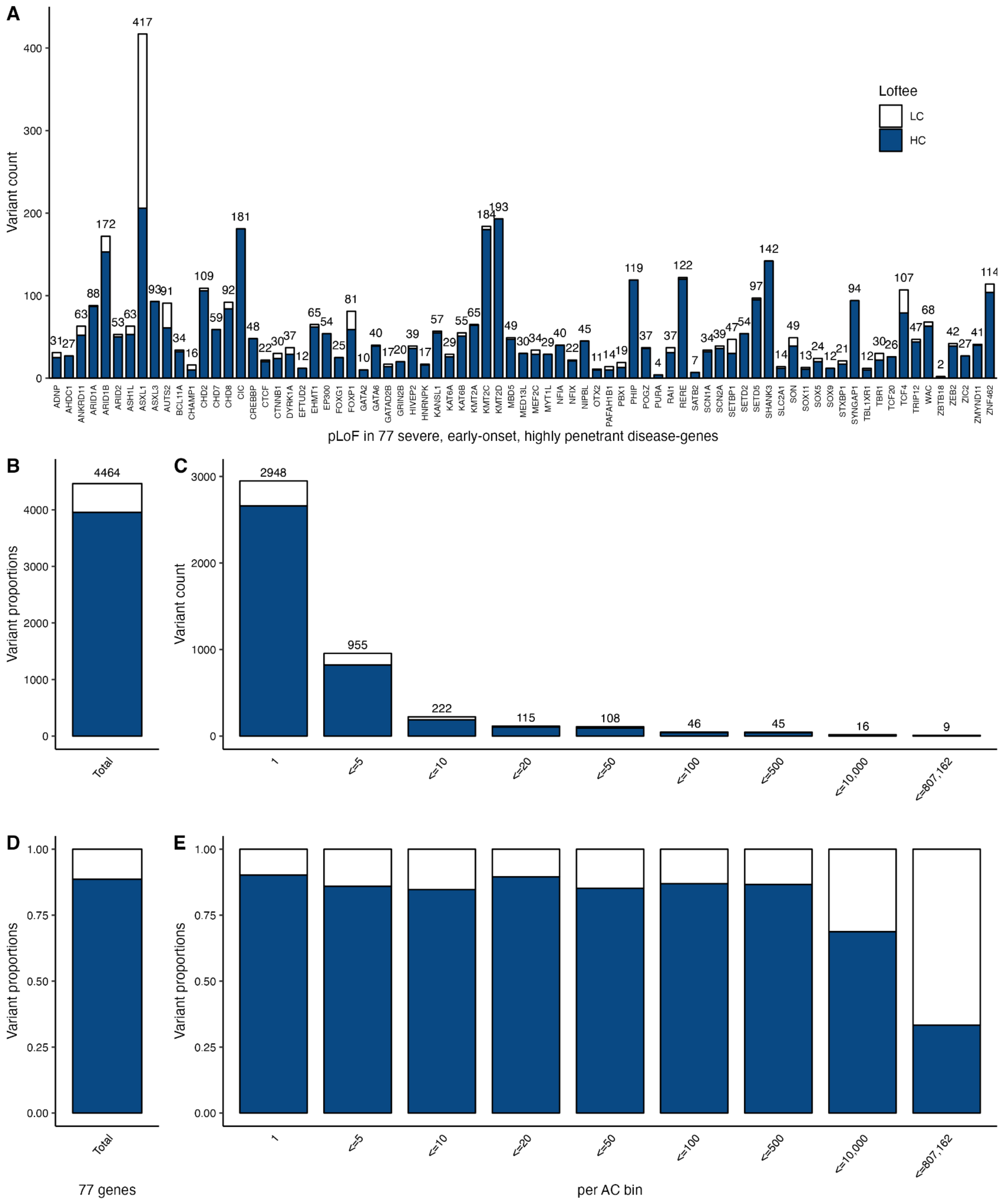


**Fig. S4:** pLoF variants in the 77 genes in 807,162 individuals in gnomAD v4. (A) Variant count per gene, colored by low confidence (LC, white) or high-confidence (HC, blue) by Loss-Of-Function Transcript Effect Estimator (LOFTEE). (B) The total number of variants and (C) variants per allele count (AC) bin. (D-E) Proportion of LC and HC variants for all variants (D), per AC bin (E).

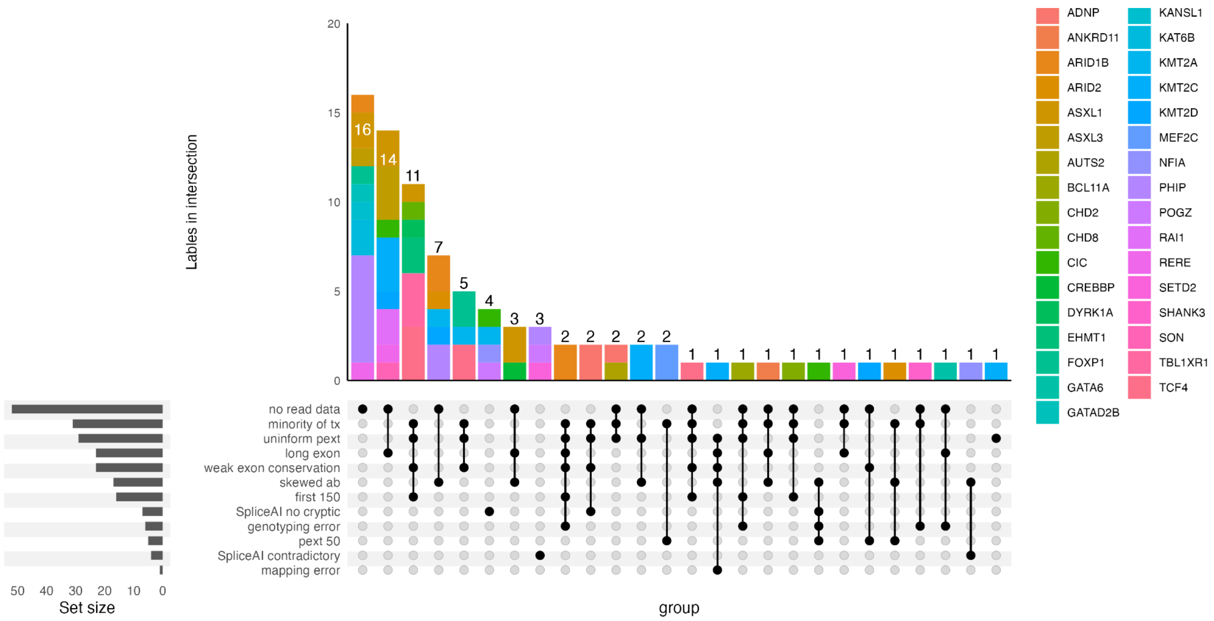


**Fig. S5:** Upsetplot for variants labeled as uncertain, colored by gene.

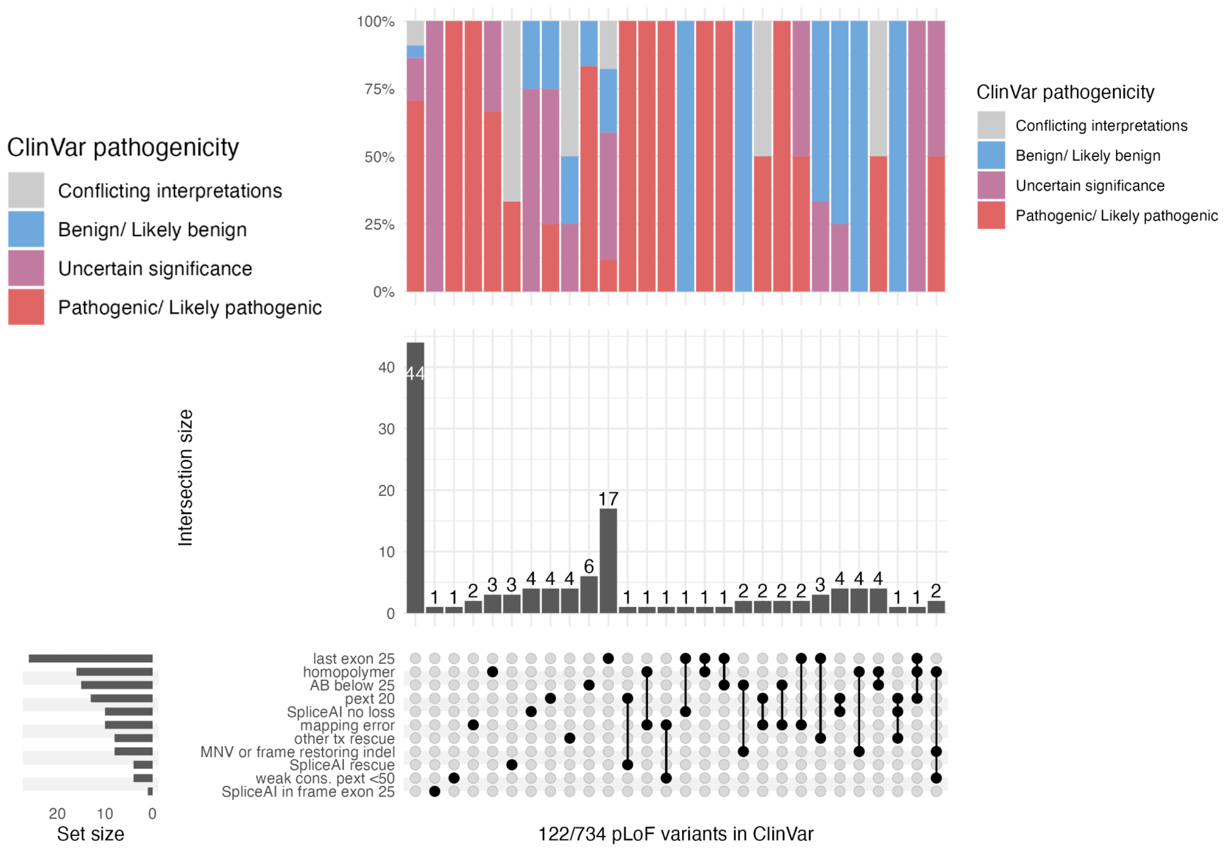


**Fig. S6:** Upsetplot of 122 of 734 pLoF also reported in ClinVar including labels that result in a not/likely not pLoF verdict listed, colored by ClinVar pathogenicity category. The 44 variants (first bar) with no flags had a verdict of “LoF”, “Likely LoF” or “Uncertain”.

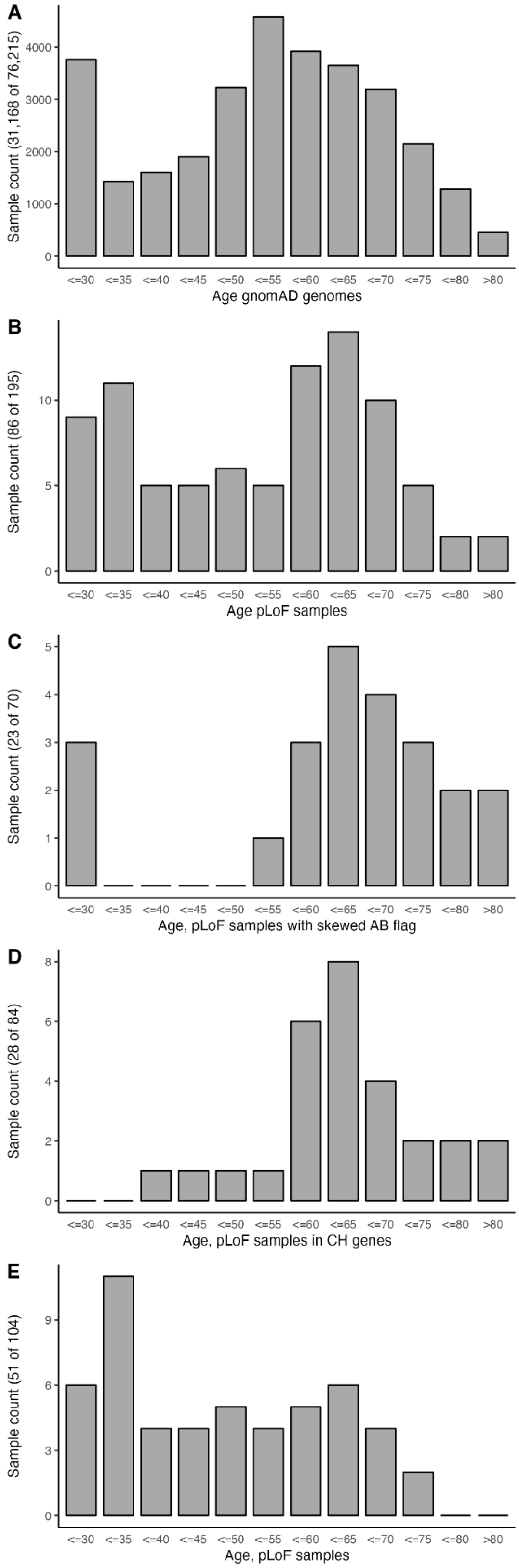


**Fig. S7:** Skewed age and allele balance (AB) as an indicator of somatic variation. (A) Age distribution in gnomAD genomes, aga data available for 31,168 of 76,215 individuals. (B) Age distribution in 195 samples with 104 pLoF variants passing LoF curation and excluding pLoF in samples from the 1000 Genomes Project, age data available for 86 of 195 samples. (C) Age distribution in 70 samples where the pLoF shows skewed AB ratio (alternative allele >25% but <35%), age data available for 23 of 70 samples. (D) Age distribution in 84 samples with pLoF variant in genes associated with clonal hematopoiesis (CH), age data available for 28 of 84 samples. (E) Age distribution of remaining 104 samples with 50 pLoF variants passing somatic filtering, age data available for 51 of 104 samples.

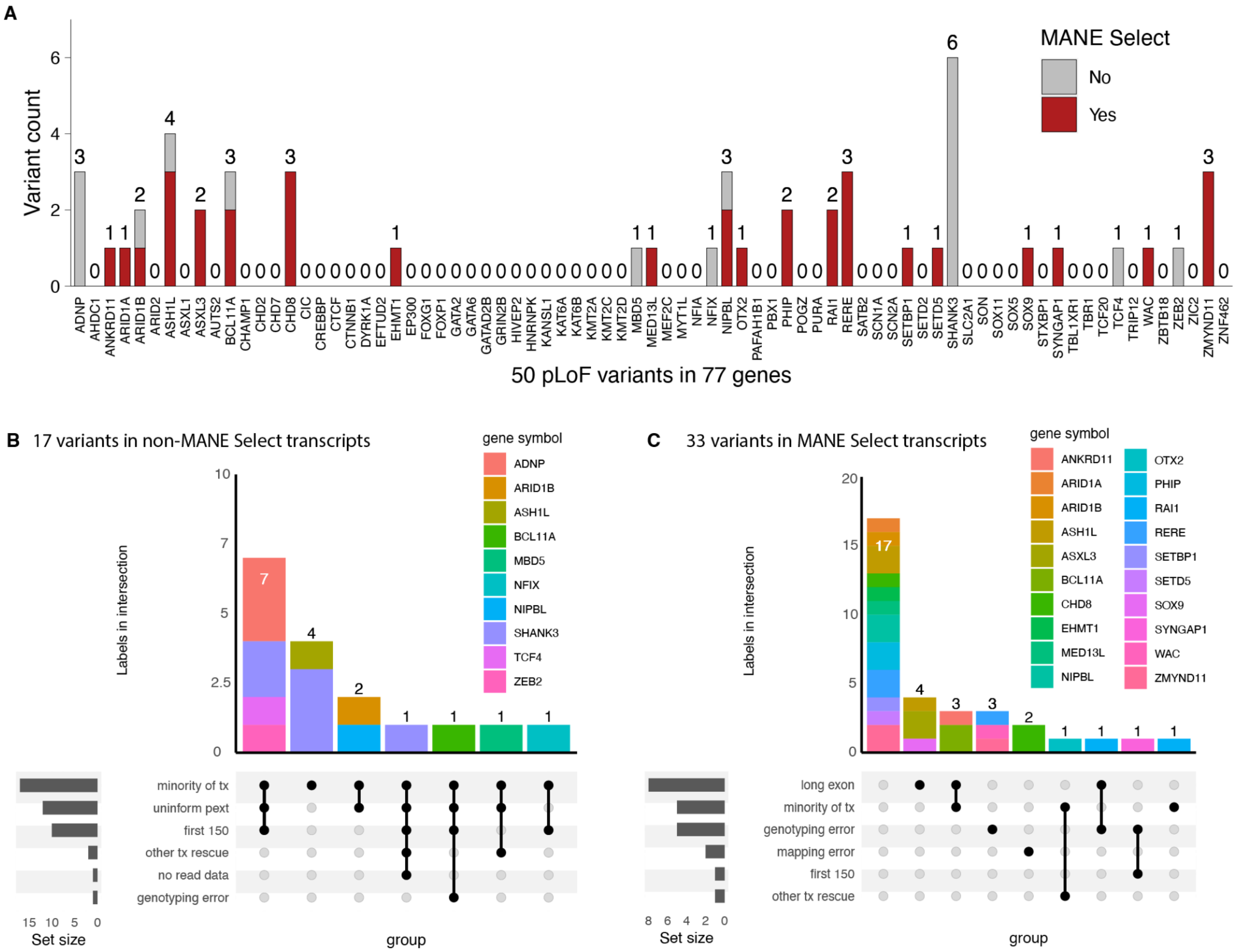


**Fig. S8:** Of 734 pLoF variants in 77 genes associated with haploinsufficiency disease, only 50 pLoF variants (6.8%) remained unexplained after careful scrutinization (A) Variant count per gene colored by whether the variant is present in the MANE Select transcript (red) or not (grey). (B) Upset plot of features of 17 variants in non-MANE Select transcripts and (C) 33 pLoF variants in MANE Select transcripts.

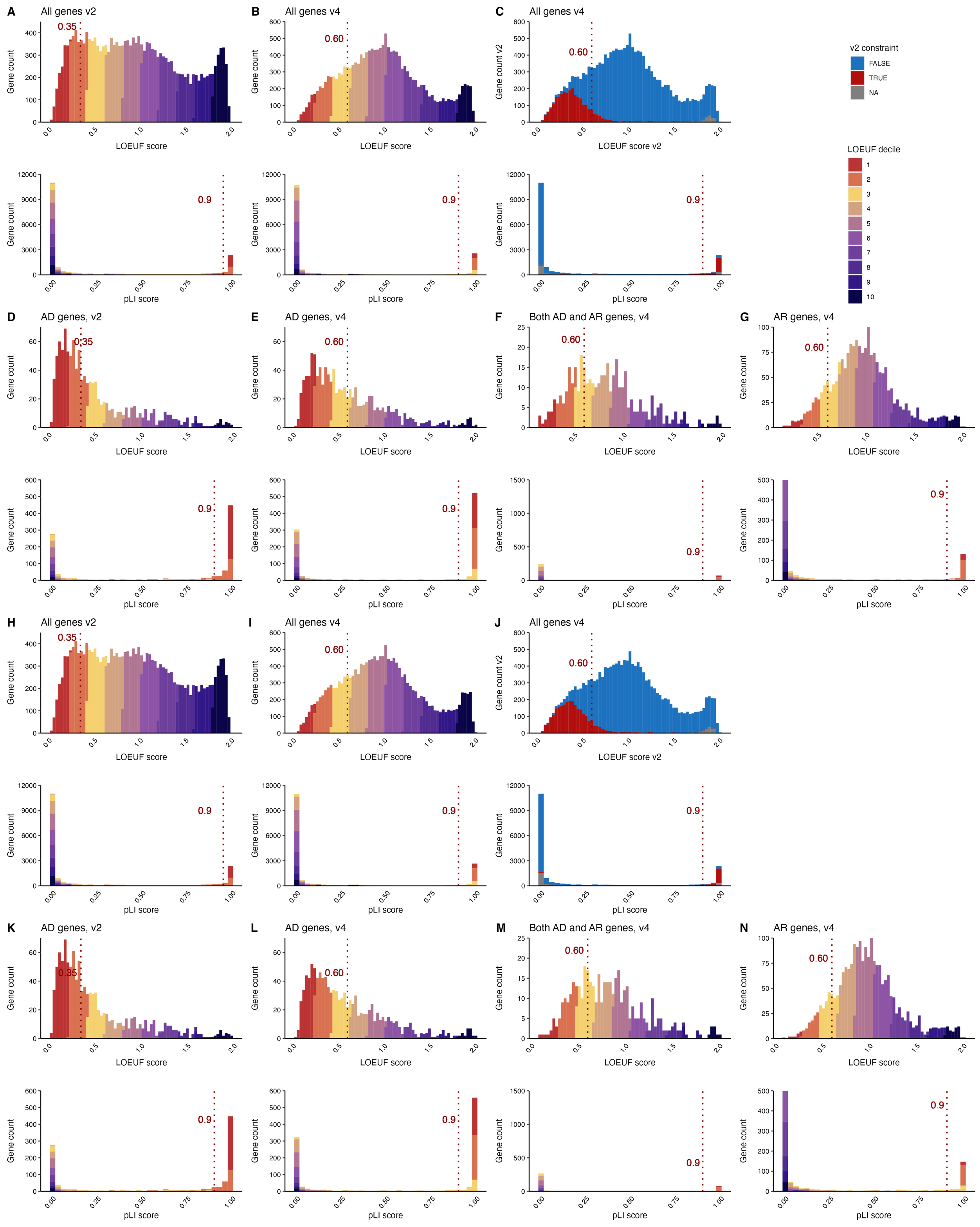


**Fig. S9:** Constraint scores, LOEUF score (upper) and pLI (lower), colored by LOEUF decile (A) for all genes in v2, (B) all genes in v4, (C) constraint scores in v4 colored by constraint in v2

(LOEUF ≤ 0.35, pLI≥  0.9, red) and not constraint in v2 (LOEUF >0.35, pLI <0.9, blue). (D) Distribution of constraint scores for genes associated with diseases of autosomal dominant (AD) in v2 (D) and v4 (E), as well as for v4 both AD and autosomal recessive (AR), and (F) AR inheritance in OMIM (G). Transcripts for v4 match transcripts of v2. Dotted line indicates threshold for loss-of-function constraint (0.35 in v2 and 0.60 in v4 for LOEUF, 0.9 for pLI). (H)-(N) Replication of A-G using MANE Select transcripts in v4 (rather than matching v4 transcripts to v2 in A-G).

### Supplementary Tables

**Table legends**

**Table S1:** Allele count for allele frequency (AF) bins, Fig. 1d.

**Table S2:** Allele count for P/LP allele count (AC) bins, Fig. 1e.

**Table S3:** Assessment of 90 unique P/LP occurring in combination with pLoF when investigating local pLoF as rescue of 3957 P/LP in ClinVar variants.

**Table S4:** 77 haploinsufficiency genes associated with severe disorders not expected to be compatible with participation in common disease studies (e.g., gnomAD), early onset (before the age of three), and reported as highly penetrant (using *de novo* rate as a proxy for penetrance).

**Table S5:** The specific set of rules used for this project modified for conservative curation using the advanced framework for loss-of-function curation previously published by this group[25]**.**

**Table S6:** Full curation results of 734 pLoF variants found in 77 haploinsufficiency genes in gnomAD v4 genomes.

**Table S7:** Counts for each ClinVar classification and pLoF curation category, ClinVar data was available for 122 of 734 pLoF variants (in Figure 3g).

**Table S8:** sQTL results from all pLoF variants.

**Table S9:** Genotypes for sQTL in individuals with MEF2C pLoF variants.
